# Age- and Sex-Related Differences in Sleep Patterns and Their Relations to Self-Reported Sleep and Mood

**DOI:** 10.1101/2025.05.15.654262

**Authors:** Habiballah Rahimi-Eichi, Justin T. Baker, Anders M. Fjell, Randy L. Buckner

**Affiliations:** Department of Psychology, Center for Brain Science, Harvard University, Cambridge, MA 02138; Institute for Technology in Psychiatry, McLean Hospital, Belmont, MA 02478; Department of Psychiatry, Harvard Medical School, Boston, MA 02114; Center for Lifespan Changes in Brain and Cognition, University of Oslo, Oslo, Norway; Center for Computational Radiology and Artificial Intelligence, Oslo University Hospital, Oslo, Norway; Athinoula A. Martinos Center for Biomedical Imaging, Massachusetts General Hospital, Boston, Charlestown, MA 02129

**Keywords:** UK Biobank, Aging, Actigraphy, Depression

## Abstract

Robust age- and sex-related differences in sleep were identified (n=38,546) and replicated (n=38,547) from week-long passive actigraphy data in UK Biobank participants ages 44-82. Sleep patterns reflected reliable non-linear interactions between age and sex. Younger women slept about 17 min more than their male counterparts, though this difference diminished with age, with both sexes reducing total sleep duration in later life. Middle-aged individuals exhibited shorter sleep durations during the week, with weekend sleep increasing by as much as 50 min. Participants in their seventh and eighth decades showed more consistent sleep patterns throughout the week. Sleep patterns also suggest maintenance of total sleep duration: individuals reporting waking too early maintain sleep duration by going to sleep earlier, while individuals reporting sleeping too much fall asleep later but also wake later, again maintaining sleep duration. Self-reported depression and anhedonia were associated with reduced total sleep duration across multiple age groups and both sexes. These collective results indicate that the timing, consistency, and overall amount of sleep differs by age and is affected by individual factors.

## Introduction

Sleep is an essential biological function that supports diverse physiological and psychological processes, including metabolic health, immune system resilience, and cognitive performance (Capone *et al*., 2019; Bar *et al*., 2021). Disruptions in sleep patterns, such as reduced sleep efficiency, early morning awakenings, and increased wake after sleep onset (WASO), are associated with neurodegenerative diseases, including Alzheimer’s and Parkinson’s (Wang and Holtzman, 2020; Gao *et al*., 2022; Zhang *et al*., 2022). Sleep and daily activity patterns are also disrupted in mental illness suggesting relevance to a broad array of human clinical conditions (Freeman *et al*., 2020; Scott *et al*., 2021). It is thus critical to understand how sleep patterns differ by age and sex, and how patterns interact with mood and well-being. Here we provide such a normative reference by estimating sleep patterns under natural conditions using objective actigraphy in a large cohort of 77,093 individuals.

While previous research has examined sleep patterns across age and sex, the reliance on self-reported data can introduce bias, particularly among older adults who may have altered self-perceptions of sleep (Schoeler, Pingault and Kutalik, 2024; Jackson *et al*., 2020; Van den Berg *et al*., 2008). To address these limitations, objective sleep tracking methods, such as actigraphy, are increasingly being used for large-scale, real-world studies. Although polysomnography (PSG) remains the gold standard, it is often impractical for continuous, population-wide monitoring. Actigraphy-based measures of sleep provide objective assessment of sleep and activity in natural settings at an unprecedented scale (van Hees *et al*., 2015; Windred *et al*., 2021).

In this study, we examined daily sleep and wake activity in middle-aged and older adults (ages 44–82) using week-long actigraphy data from the UK Biobank. Through rigorous quality control measures, we ensured the inclusion of only participants with complete, continuous data over six days, allowing us to capture robust sleep parameters. Our analyses explored sleep duration, onset, and wake time, with a particular focus on age- and sex-related differences. Additionally, we investigated the relationship between objective sleep parameters and self-reported measures to understand how self-reported perceptions of sleep link to objective estimates, as well as to understand how mental health symptoms, particularly depression and anhedonia, influence objective sleep measures. Our analyses revealed robust interactions between sleep, aging, and mental well-being with unexpected and sometimes counterintuitive patterns.

## Materials and Methods

### Participants

Participants were drawn from the UK Biobank, a large-scale cohort of over 500,000 individuals at recruitment, who were assessed at 22 centers across the United Kingdom (Sudlow *et al., 2015)*. This cohort provides a semi- representative sample of the population, encompassing a wide range of sociodemographic, lifestyle, and health characteristics (Fry *et al., 2017*). A subset of 111,635 participants, ages 44–82 years, was randomly selected during 2014–2015 to wear wrist-worn accelerometers for the passive measurement of physical activity levels, after providing informed consent (Doherty *et al*., 2017). In 2017, an online follow-up questionnaire focused on mental health, largely based on the World Health Organization (WHO) Composite International Diagnostic Interview (CIDI), was completed by approximately 159,000 participants (Davis *et al*., 2020). From this assessment, questions related to sleep, depression, and anhedonia were utilized here to compare self-reports of sleep and mood with objective activity measures. Ethical approval for data re-analysis was granted by the Harvard University Human Studies Committee under UK Biobank Application 67237.

### Estimation of Activity Levels and Major Sleep Episode from Raw Accelerometer Data

Accelerometer data were obtained using a 3-axis wristband device (model AX3; Axivity Ltd, Newcastle, UK), which recorded participants’ movements continuously for up to 8 days (Doherty *et al*., 2017). To derive objective sleep and activity metrics, we applied the open-source DPSleep pipeline (Rahimi-Eichi *et al*., 2021), modified to handle UK Biobank data (Figure 1). This pipeline utilized the standard deviation of accelerometer signals to detect wrist-off episodes and calculated minute-level power across the 3 axes using power density spectrum analysis. Within-individual activity levels were classified by percentiles (10th, 25th, 50th, and 75^th^) using iterative application of sliding windows (100, 60, and 90 minutes) to robustly estimate the major sleep episode (Rahimi-Eichi *et al*., 2021). The longest period of low activity was designated as the sleep episode, without prior assumptions about sleep onset time or duration. Automatic hierarchical rules were applied to refine sleep episode boundaries and merge short adjacent sleep bouts, yielding the final estimate of the major sleep episode. Further details on the validation and limitations of these estimates can be found in Rahimi-Eichi et al. (2021).

**Figure 1.**
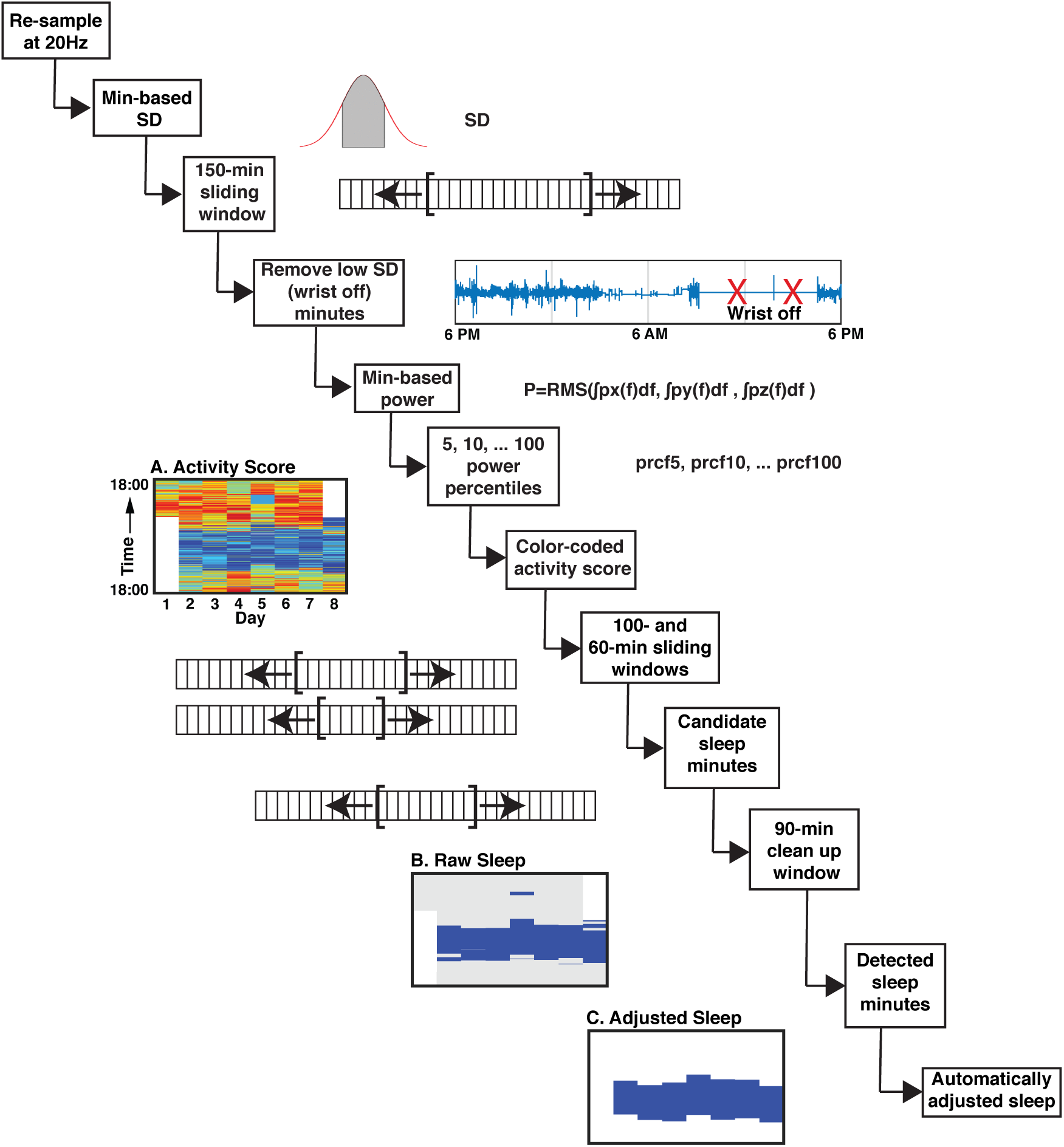
Processing pipeline used to estimate sleep duration. The sequential steps of the fully automated processing pipeline used to estimate the major sleep epoch from the UK Biobank data are illustrated. The pipeline, referred to as DPSleep (Rahimi-Eichi et al. 2021), begins with resampling raw accelerometer data, generating individualized activity scores (A), raw (B), and adjusted sleep episodes (C). Sleep parameters are determined based on automatically adjusted sleep episodes as shown in panel C. The raw data are resampled at 20Hz and the standard deviations of acceleration across the 3 axes are averaged over 150-min intervals forward and backward to identify wrist-off minutes. The power density spectrum of the acceleration signal is then computed, and the area under the curve approximates the raw activity power as the root mean square integrated across the axes. Every minute is categorized into percentile thresholds, from 5 to 100, for visualization (A; dark blue to red indicates increasing activity scores) and quantification. Specific thresholds (i.e., 10, 25, 50 and 75 percentiles) are used to determine sleep episodes with a series of forward and backward moving averages applied to identify potential sleep episodes, which are then refined to produce the raw sleep estimate (B) and yield the final adjusted sleep estimate (C).

### Accelerometer Data Quality Control (QC)

Accelerometer data from the UK Biobank were first screened to exclude duplicate samples from the same individual, as shown in Figure 2. Remaining data were processed using the DPSleep pipeline. After processing, datasets were further excluded if they exhibited structural anomalies or had more than four days of missing sleep data. A second round of manual quality control (QC) was then conducted, as depicted in Figure 3. This QC process was blinded to participants’ age and sex. Condensed activity plots, displaying daily activity patterns from 200 participants each, were reviewed for missing data or off-wrist periods (Figure 3). Off-wrist periods were allowed only on days 1 and 8, provided they did not interfere with the measurement of sleep onset on day 2 or wake time on day 7 ensuring six continuous days of data. Participants whose data coincided with daylight-saving time changes were excluded. These comprehensive QC procedures ensured inclusion of only high-quality data with sufficient sampling for accurate sleep pattern estimation.

**Figure 2.**
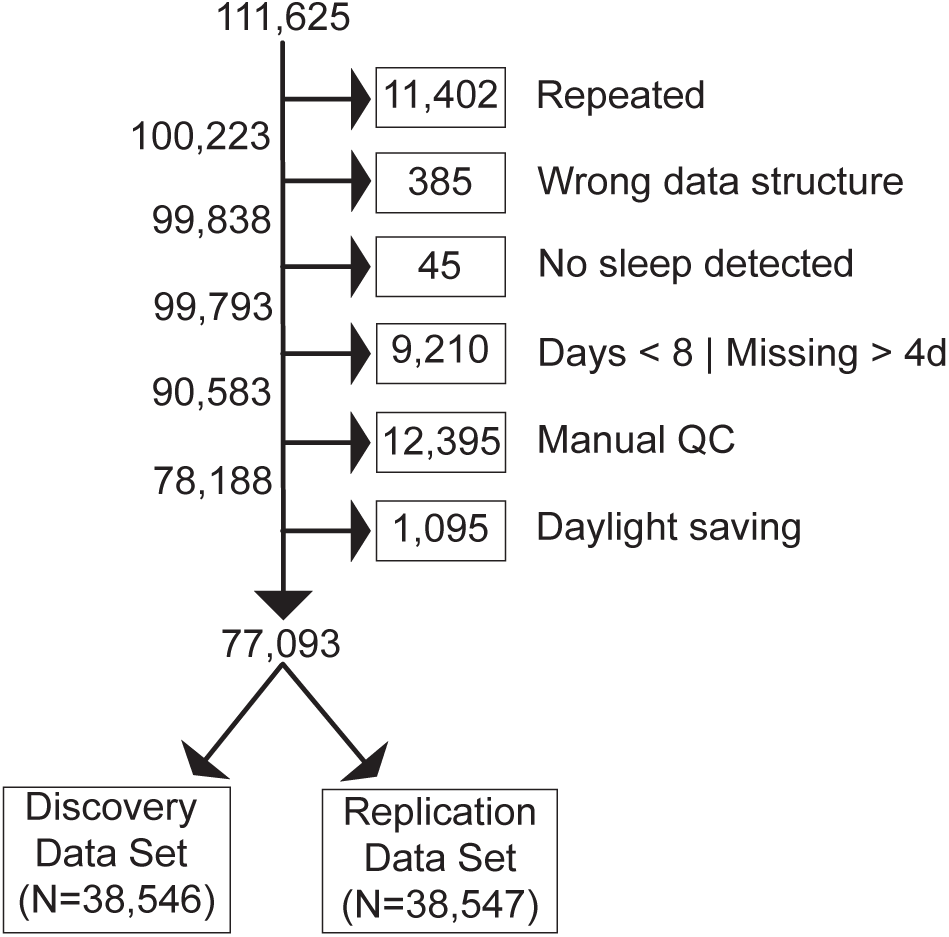
Flow chart of participant inclusion. A flow chart illustrates the participant selection that resulted in two independent Discovery and Replication datasets used for all subsequent analyses. Out of 111,625 initial candidate samples exclusions included repeat samples, wrong data structure, no sleep detected, missing data, manual QC failures (see Figure 3), and exclusion for daylight savings time. These exclusions were all made before examining sleep structure to mitigate bias. The final sample included 77,093 participants, divided into two equal groups for Discovery (N=38,546) and Replication (N=38,547). To visualize effects of age and sex the data were further separated into age bins (44-49, 50-54, 55-59, 60-64, 65-69, 70-74, 75-82) and separated groups by genetic sex (XX = genetic female; XY = genetic male).

**Figure 3.**
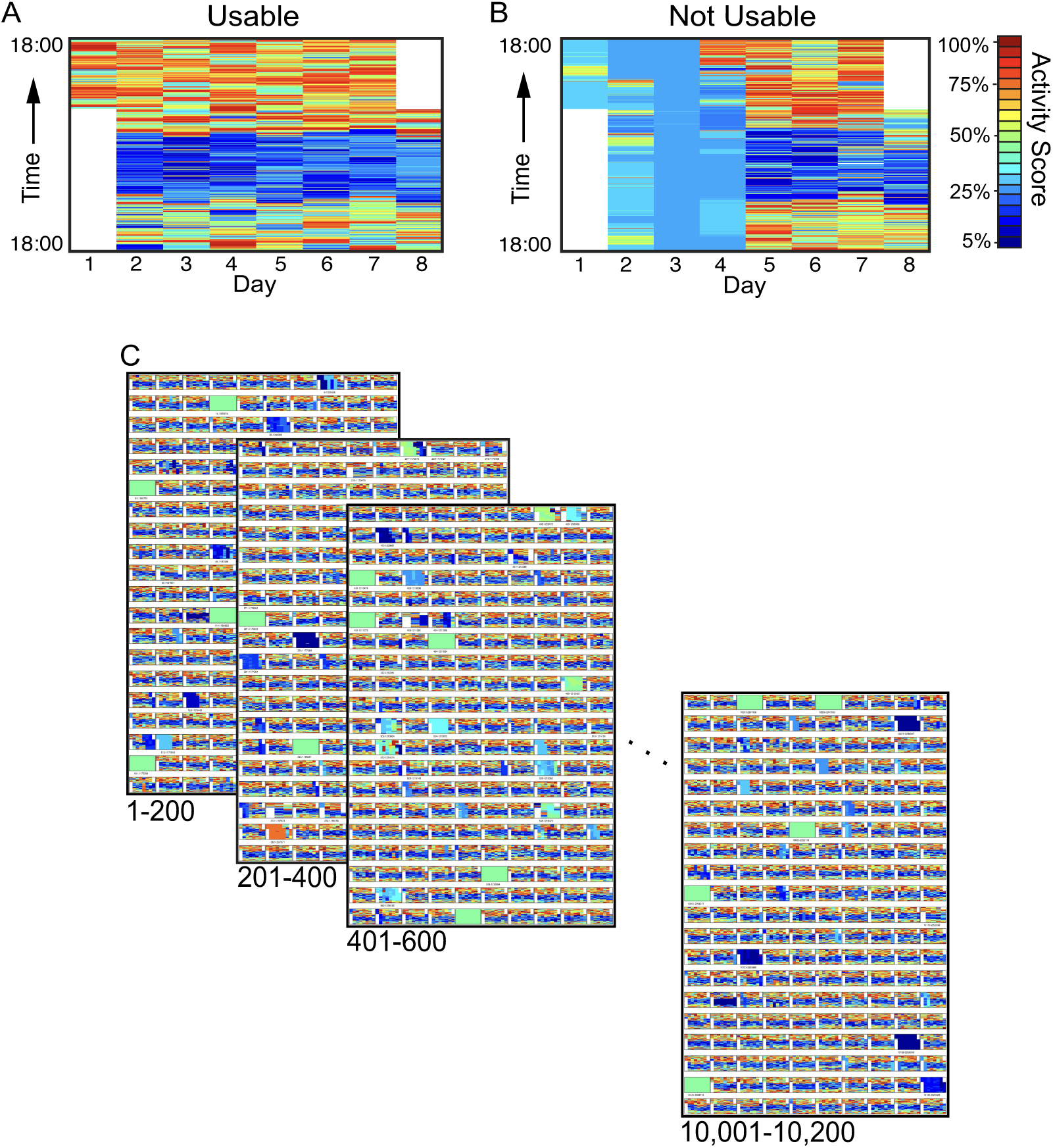
Manual quality control. To ensure only high-quality data was included in subsequent analyses, a manual quality control (QC) procedure was developed that could efficiently accommodate the large sample size. Exclusions were made blind to age or any other features of the data to minimize bias. Every sample was visualized as a week-long panel that showed the day-by-day estimated activity. A usable example (A) with complete actigraphy data for 6 consecutive nights and days is shown alongside an unusable example (B) with several missing days, indicated by flat or low-variance activity scores (blue). QC maps (C) display activity scores created for 200 participants per page, allowing review of every sample. Only individuals with complete sleep and wake accelerometer data for days 2 though 7 were included in the analysis (days 1 and 8 were often truncated, as expected, reflecting watch wearing began and ended).

The final resulting sample of 77,093 participants was divided into two equally sized datasets: Discovery (N = 38,546) and Replication (N = 38,547) (Figure 2). These datasets were matched by sex and grouped into age bins of 44–49, 50–54, 55–59, 60–64, 65–69, 70–74, and 75–82 years (Table I). The separation into two datasets allowed for independent replication of findings.

**Table I.** Participant counts after QC stratified by age group and sex (N=77,093)

|  | **44-49** | **50-54** | **55-59** | **60-64** | **65-69** | **70-74** | **75-82** | **All Ages** |
| --- | --- | --- | --- | --- | --- | --- | --- | --- |
| **XX** | 2,904<br>(3.77%) | 5,909<br>(7.66%) | 7,016<br>(9.10%) | 8,760<br>(11.36%) | 10,737<br>(13.93%) | 6,829<br>(8.86%) | 1,249<br>(1.62%) | 43,404<br>(56.30%) |
| **XY** | 1,995<br>(2.59%) | 3,854<br>(5.00%) | 4,411<br>(5.72%) | 6,124<br>(7.94%) | 8,952<br>(11.61%) | 6,912<br>(8.97%) | 1,441<br>(1.87%) | 33,689<br>(43.70%) |

### Self-Reported Sleep and Mood

The UK Biobank online questionnaires included several items assessing mental health and well-being (Davis *et al*., 2020). Participants were asked whether their sleep had changed, with a binary response option ("Yes" or "No") to the question, "Did your sleep change?" Follow-up questions on "sleeping too much" (Data-Field 20534) and "waking too early" (Data-Field 20535) were analyzed to investigate associations between self-reported sleep disturbances and objective sleep measurements. Additionally, during the baseline assessment, participants were asked about their sleep duration through the question, "How many hours of sleep do you get in every 24 hours (please include naps)?" (Data-Field 1160) (Sudlow *et al*., 2015). Participants could select any integer to respond. We analyzed the association between self-reported sleep duration and objective sleep measurements to ensure consistency and validity.

In the section on depression, participants rated the severity of their symptoms over the past two weeks, including feelings of depression (Data-Field 20510) and anhedonia (lack of pleasure) (Data-Field 20514). Response options included: "not at all," "several days," "more than half the days," and "nearly every day." For analysis, responses were dichotomized into two categories: "No" for "not at all," and "Yes" for any response indicating symptoms for "several days," "more than half the days," or "nearly every day."

### Statistical Analysis

The mean and standard error (SE) were calculated to describe the effects of age, sex, and day of the week on sleep parameters. Comparison of self-reported sleep duration groups was conducted using one-way ANOVA; binary self-report groups (i.e., "Yes" vs. "No") were compared using two-sample t-tests, with significance set at *p* < 0.05 for both tests. In the instances of multiple groups, one-way ANOVAs were utilized with significance set at *p* < 0.05. Statistical analyses were performed using MATLAB R2022b with the Statistics and Machine Learning Toolbox (MathWorks, Natick, MA, USA).

## Results

The results provide a comprehensive description of how objective sleep and activity measures vary by age and sex, and illuminate associations between these measures and self-reported sleep quality and mental health symptoms. To ensure that findings are robust, key analyses were replicated in independent Discovery and Replication datasets. Note also that each age and sex group (within each dataset) is itself an independent sample. Thus, when multiple tests are made across groups or replication datasets that converge on a pattern, these should be viewed as replications in independents samples, as they are not multiple tests on the sample same. That is, they illustrate generalizations across datasets.

### Sleep Patterns Differ by Age and Sex

Replicable differences were observed in sleep duration, sleep onset, and wake time when examined in relation to age and sex. On average, men exhibited shorter sleep durations compared to women across most age groups, particularly in those under 60 (Figure 4). This disparity was driven by differences in sleep onset and wake times, with men going to bed later and waking earlier than women. Specifically, men under 60 went to bed later than women, yet woke up earlier, leading to about 17 min in reduced total sleep duration (Figure 4, Table II).

**Figure 4.**
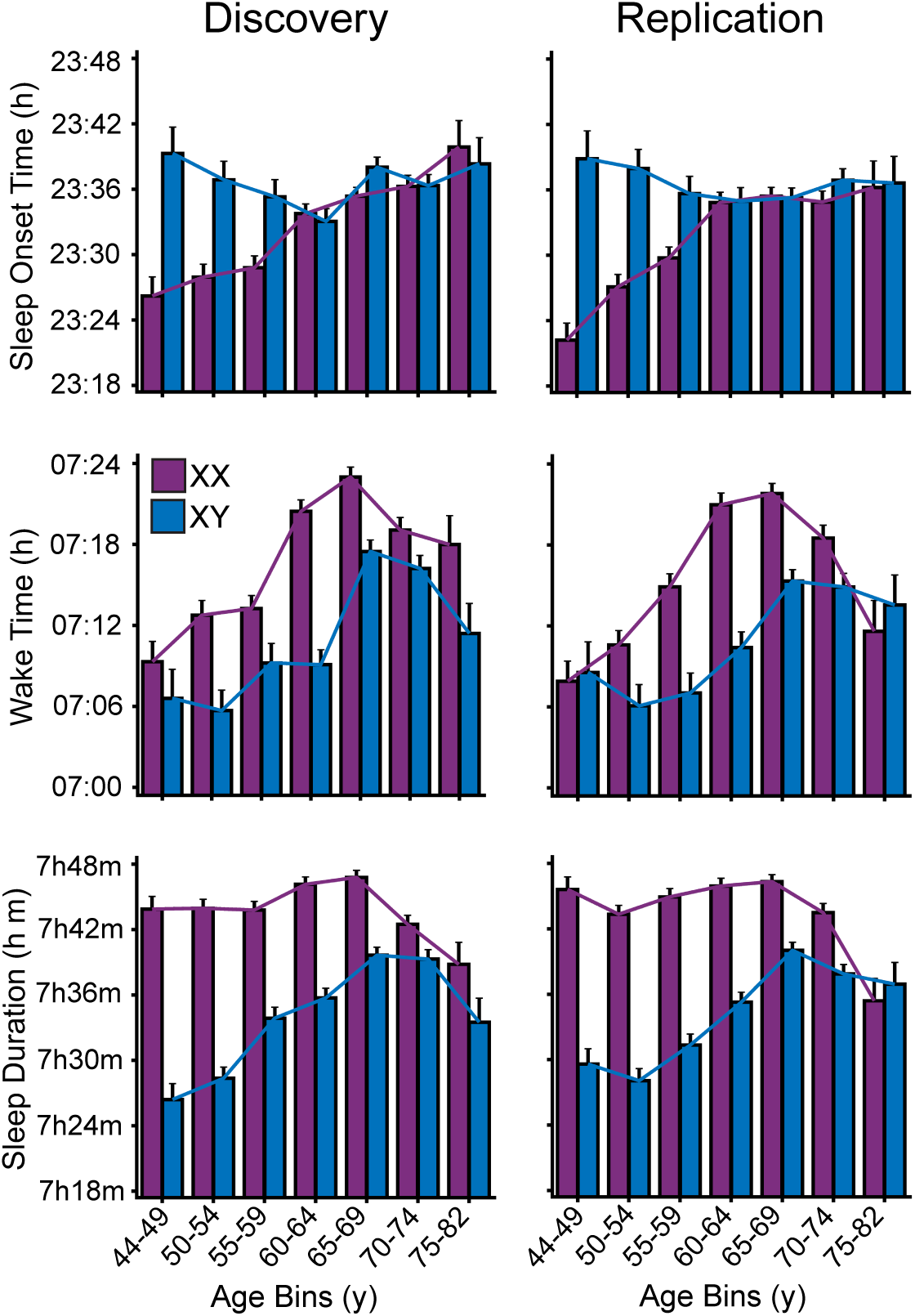
Sleep patterns differ by age and sex. Sleep onset, wake time, and sleep duration are shown for separate age and sex groups. Mean and standard error are displayed for each parameter for each group, where each bar is a distinct group of participants. The left panels display the sleep patterns for the Discovery dataset and the right panels the patterns for the independent Replication dataset. Men younger than 60 generally go to sleep later and wake up earlier than women, resulting in shorter sleep duration by up to 20 min. This sex-associated sleep gap diminishes with age. Older men’s sleep durations are longer and older women’s shorter. Older age groups for both sexes tend to wake up later until the 70s, where this trend plateaus or reverses. These idiosyncratic age by sex interaction patterns are robust and replicate across the Discovery and independent Replication datasets. h = hour; y = year.

**Table II.** Independent Samples t-Test Results for Sleep Parameters Between Sexes.

|  | **44-49** | **50-54** | **55-59** | **60-64** | **65-69** | **70-74** | **75-82** |
| --- | --- | --- | --- | --- | --- | --- | --- |
| **Sleep Onset Time** | t(4782) = -5.46<br>p<0.001 | t(8992) = -5.80<br>p<0.001 | t(10746) = -3.14<br>p<0.01 | t(14281) = 0.18 | t(19306) = -1.44 | t(13630) = -0.89 | t(4782) = -0.81 |
| **Wake Time** | t(4782) = 3.54<br>p<0.001 | t(8992) = 5.32<br>p<0.001 | t(10746) = 6.61<br>p<0.001 | t(14281) = 10.56<br>p<0.001 | t(19306) = 6.58<br>p<0.001 | t(13630) = 3.27<br>p<0.01 | t(4782) = 1.70<br>p<0.1 |
| **Sleep Duration** | t(4782) = 12.52<br>p<0.001 | t(8992) = 15.04<br>p<0.001 | t(10746) = 12.52<br>p<0.001 | t(14281) = 12.64<br>p<0.001 | t(19306) = 9.37<br>p<0.001 | t(13630) = 4.77<br>p<0.001 | t(4782) = 0.90 |

**Table III.** Independent Samples t-Test Results for Wake Activity Between Sexes.

|  | **44-49** | **50-54** | **55-59** | **60-64** | **65-69** | **70-74** | **75-82** |
| --- | --- | --- | --- | --- | --- | --- | --- |
| **Wake Activity** | t(4782) = 0.28 | t(8992) = -0.46 | t(10746) = -0.36 | t(14281) = 4.82<br>p<0.001 | t(19306) = 9.32<br>p<0.001 | t(13630) = 10.82<br>p<0.001 | t(4782) = 5.46<br>p<0.001 |

In the oldest participant groups, the gap between men and women diminished. By the seventh decade of life, men and women demonstrated remarkably similar sleep durations, and both sexes showed earlier wake times. This pattern may suggest that the influence of social or occupational factors on sleep duration lessens with age. Given these are cross-sectional group difference estimates, another plausible explanation for this pattern is the higher mortality rate among men in middle age, which could result in the surviving older male sample exhibiting sleep traits more akin to those of women.

Furthermore, in both men and women, the overall trend showed that individuals tended to wake later during their 60s, a trend that stabilized or reversed with increasing age. This observation could reflect a biological shift in sleep-wake regulation during later life stages or changes in lifestyle after retirement.

### Sleep Patterns Differ Between Weekdays and Weekends in an Age- and Sex-Dependent Manner

Participants under 60 showed marked, replicable differences in sleep timing between weekdays and weekends, with later sleep onset, wake times, and about 50 min longer sleep duration on the weekends (Figure 5). Of further interest, men displayed less variation in sleep onset between weekdays and weekends, suggesting that their late bedtimes may not always be linked to work obligations.

**Figure 5.**
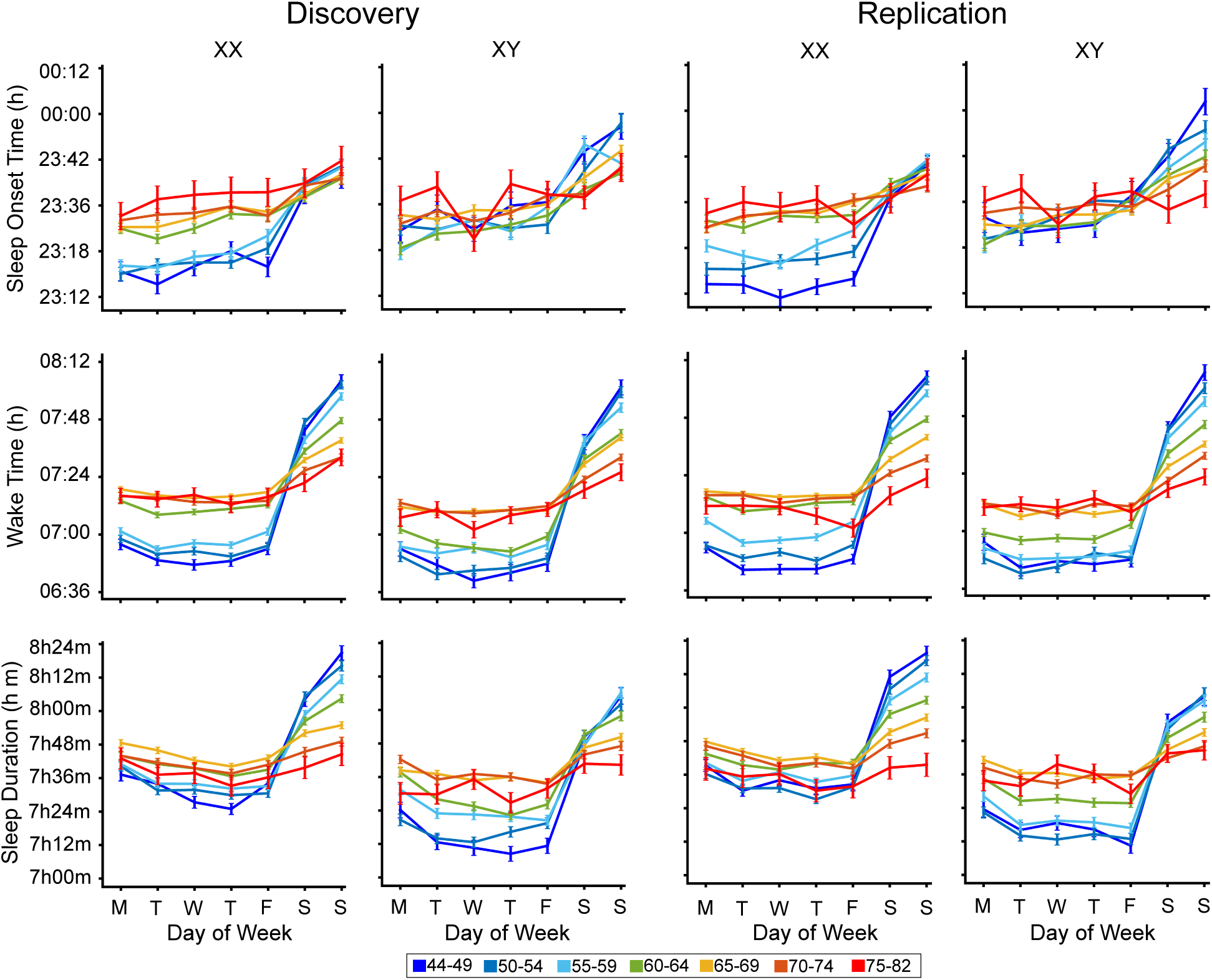
Weekly sleep patterns differ by age and sex. Sleep onset, wake time, and sleep duration are shown for each day of the week, separated into age and sex groups. Sleep parameters show significant differences between weekdays and weekends for groups under 60 years of age, with later sleep onset and wake times and longer sleep durations on weekends. Male participants younger than 60 show consistent late bedtimes across all days, while participants over 65 exhibit minimal differences between weekday and weekend sleep patterns, achieving balanced sleep duration across the week. Plots are replicated for the Discovery (left panels)) and independent Replication (right panels) datasets. XX = genetic female; XY = genetic male. M, T, W, T, F, S, S indicates day of the week beginning with Monday. Colors of the lines demarcate the age group, as illustrated by the bottom legend. h = hour.

For participants over 65, the difference between weekday and weekend sleep patterns diminished, with sleep duration and timing remaining consistent throughout the week. This uniformity may reflect fewer external obligations such as work, which tend to regulate sleep schedules in younger populations. Whatever the origin, these complex, non-linear patterns are critical to consider in studies of sleep that contain individuals of different ages.

### Wake Activity and Daily Activity Patterns Differ by Age and Sex

Wake activity declines with age, with a steeper reduction observed in men compared to women (Martin *et al*., 2014). Figure 6 illustrates this trend across different age groups and between sexes, as well as across weekdays and weekends. Younger participants, particularly those under 60, exhibited differences between weekdays and weekends, with higher activity reliably observed on weekends in both the independent Discovery and Replication datasets, for both sexes. In contrast, older participants displayed more stable (but lower overall) activity levels across the entire week, showing only slight variations between weekdays and weekends.

**Figure 6.**
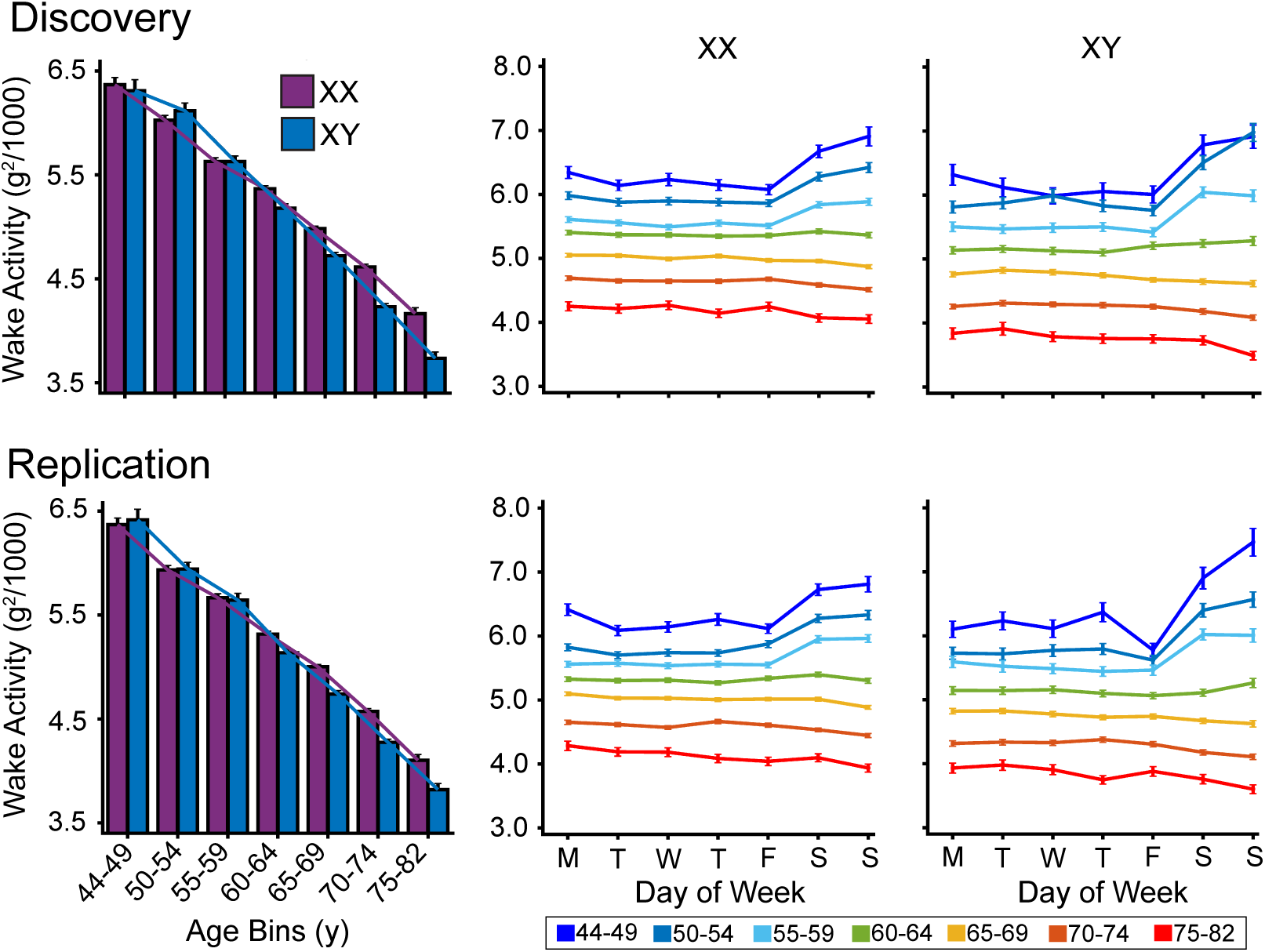
Wake activity patterns differ by age. Mean raw activity during wake episodes is shown separated by sex and age group for each day of the week. Wake activity differs (declines) significantly with age, with a slightly steeper decline observed in men. Younger participants exhibit more variation in wake activity between weekdays and weekends, particularly among men, whereas older participants maintain stable activity levels across the week. Plots are replicated for the Discovery (top panels)) and independent Replication (bottom panels) datasets. XX = genetic female; XY = genetic male. M, T, W, T, F, S, S indicates day of the week beginning with Monday. Colors of the lines demarcate the age group, as illustrated by the bottom legend.

To provide a more detailed view, Figure 7 presents the average wake activity from 6 PM of the previous day to 6 PM of the current day for all participants, mapped over the days of the week. The Discovery and Replication datasets showed consistent patterns, with higher activity in the mornings and lower activity in the evenings. This daily rhythm was evident across both samples, with a noticeable shift in sleep patterns—particularly wake times— during weekends, where mornings and afternoons were more active.

**Figure 7.**
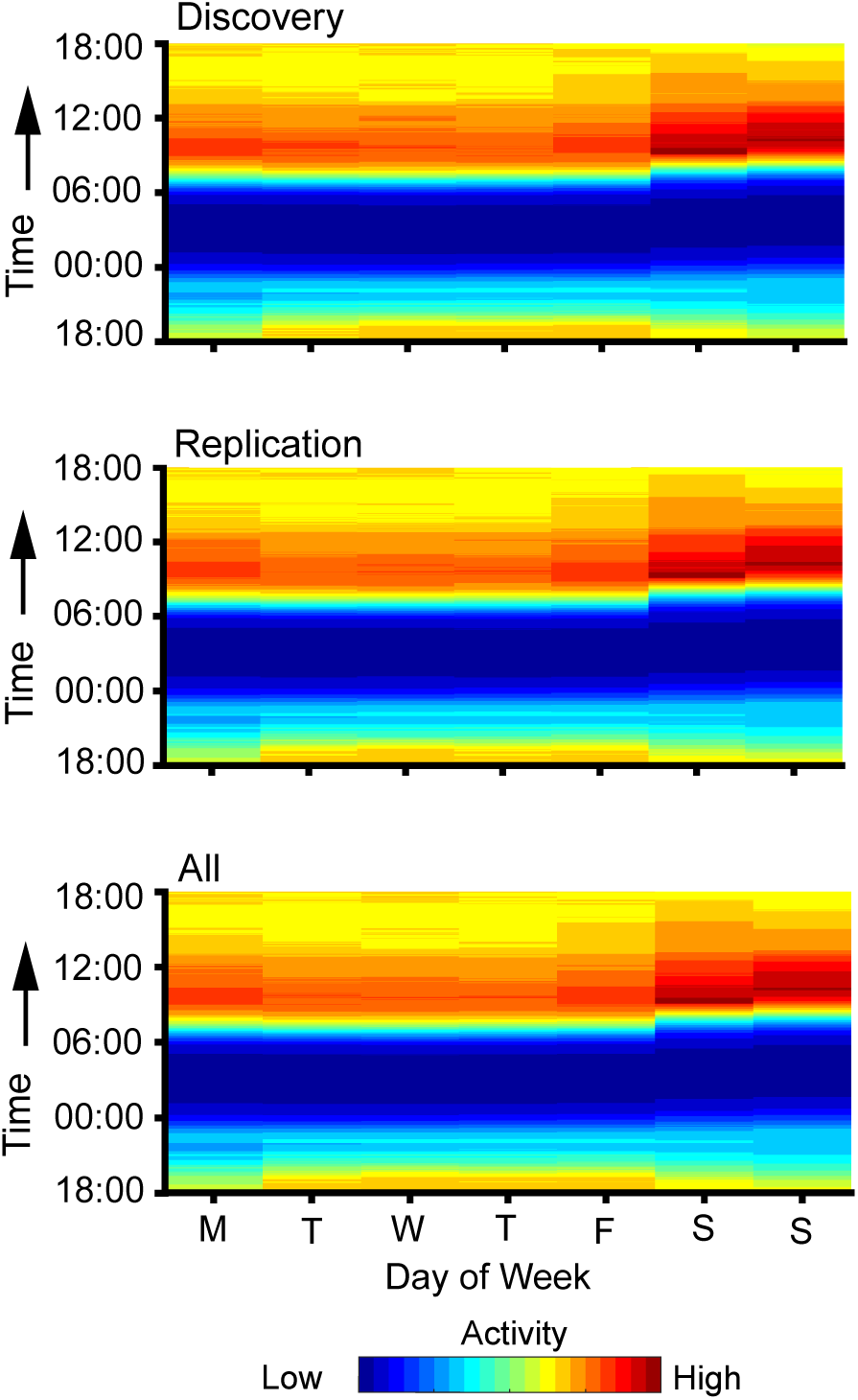
Daily activity maps split by day of the week. Mean raw daily activity from 6 PM on the previous day to 6 PM on each day is mapped across days of the week for Discovery, Replication, and the combined (All) datasets. All data are combined across age and sex. Both Discovery (top panel) and Replication (middle panel) datasets show similar patterns with higher morning activity and lower evening activity. Sleep pattern shifts, particularly in wake times during weekends, are observed consistently across samples, with more active mornings and afternoons on weekends. Colors range from dark blue to dark red, scaled to the average activity levels across all participants.

Figure 8 further breaks down the daily activity patterns by age and sex, using the fixed color scale from Figure 7. The plots illustrate a decline in activity with age in both men and women, as indicated by the reduction in higher activity (red) and the increased presence of lower activity (green). In older age groups, the distinction between activity levels during weekdays and weekends diminishes. Additionally, activity was consistently higher in the morning and lower in the evening across all age and sex groups. Notably, weekends became less active on average in the oldest participant groups.

**Figure 8.**
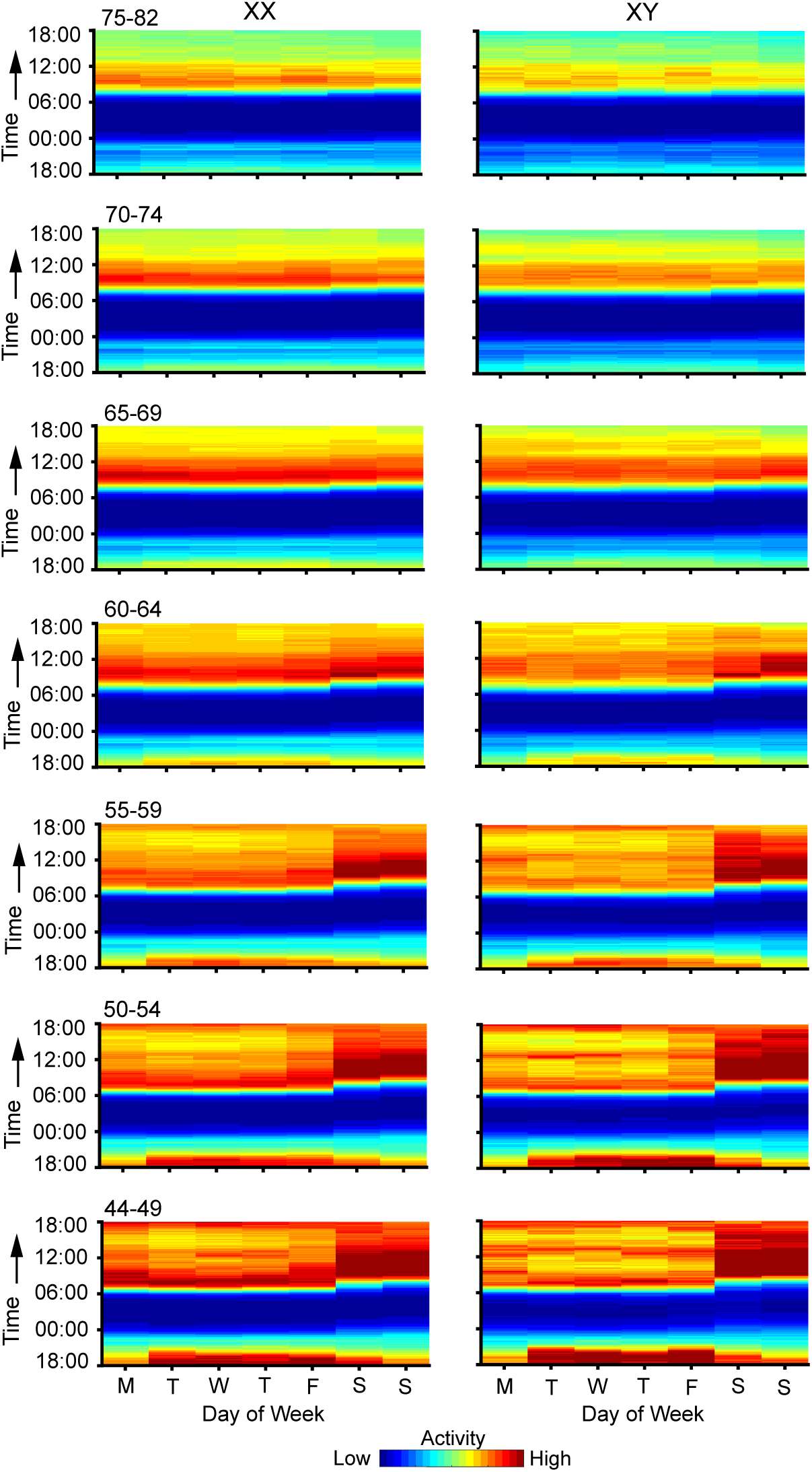
Daily activity maps split by age and sex. Mean raw daily activity for participants is displayed by sex and age group for each day of the week. The maps reveal a marked difference (decline) in activity with age for both sexes, with fewer high-activity periods (red) and more low-activity periods (green) in the oldest groups. Differences in activity levels between weekdays and weekends diminish with age. Morning activity remains higher across all groups with lower evening activity. Weekends are less active in older participants. XX = genetic female; XY = genetic male. M, T, W, T, F, S, S indicates day of the week beginning with Monday. Colors range from dark blue to dark red, scaled to the average activity levels across all participants using a fixed color scale as in Figure 7.

### Objective Sleep and Activity Differ in Relation to Self-Reported Sleep

The objective sleep measures were examined for separate samples based on self-report. As a first validity test, self-reported sleep duration was plotted in relation to objective sleep duration (Figure 9). Participants were categorized into four groups based on their self-reported sleep duration, ranging from 6 to 9 hours. Objective sleep duration was found to parametrically vary as a function of self-reported duration, consistently across age and sex groups. One-way ANOVA confirmed that the differences in objective sleep duration across the self-reported categories were independently statistically significant in many of the age groups (9 independent between-group contrasts showed a significant effect, p<0.05). In the remaining 5 between-group comparisons, the pattern was in the same direction without exception.

**Figure 9.**
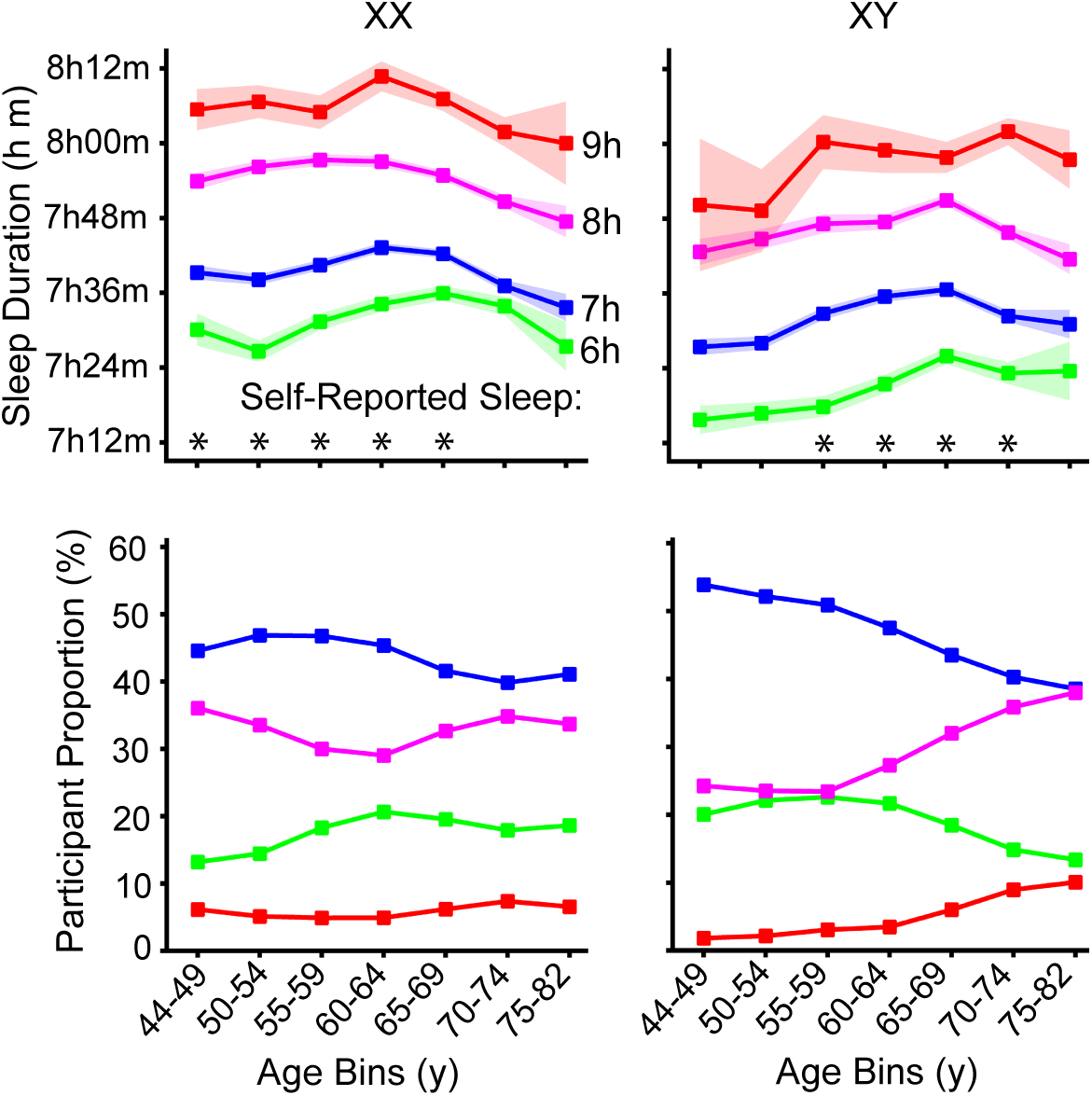
Objective sleep duration for individuals self-reporting their daily sleep duration. Objective sleep duration is shown for participants split by groups with distinct self-reported sleep duration. Colors represent different reported sleep durations: green (6 hours), blue (7 hours), magenta (8 hours), and red (9 hours). The objective sleep duration differentiates these groups across all ages for both men and women, though the range for objective sleep duration was between 7 and 8 hours (a compression relative to the self-reported duration). The proportion of participants in each age group is plotted separately for each sex and color-coded according to the corresponding self-reported sleep duration groups. Significance was tested using one-way ANOVA (* = p < 0.05).

The objective measures of sleep duration, though, did not span the same large range as the self-reported differences falling within the 7- to 8-hour range. The proportion of participants in each age group is visualized separately for each sex and color-coded by self-reported sleep duration categories. Among men, the proportion reporting longer sleep durations (i.e., 8 and 9 hours) increased with age, while the proportion reporting shorter durations (i.e., 6 hours) decreased—a trend not as pronounced in women.

The relationship between objective sleep measures and self-reported sleep disturbances, such as "too much sleep" or "waking too early," was also examined. Participants who reported "too much sleep" exhibited later sleep onset and wake times, but their total sleep duration was not significantly greater than those who did not report oversleeping (Figure 10). Interestingly, participants reporting excessive sleep tended to have lower wake activity during the day, suggesting that they may be less physically active.

**Figure 10.**
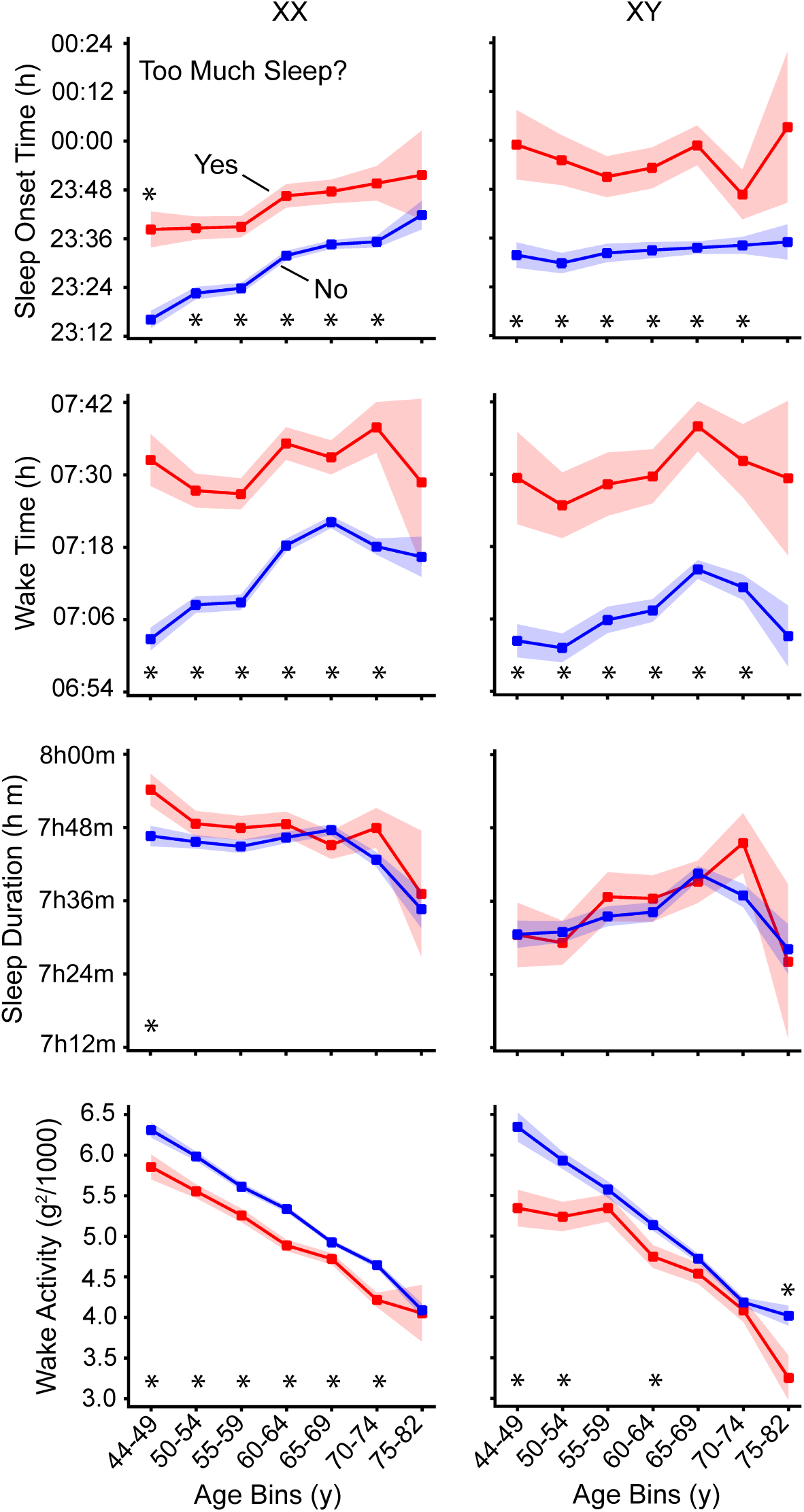
Sleep and activity parameters for individuals self-reporting "too much sleep." Sleep parameters and daily activity means are shown for participants responding "yes" or "no" to whether they were "sleeping too much." Red indicates those reporting too much sleep, and blue indicates the others, both with standard error bars. Participants reporting excess sleep wake up and go to sleep later, though their sleep duration does not significantly differ. They also show lower daily activity, possibly due to reduced overall activity or increased daytime napping. Significance was tested using a two-sample t-test (* = p < 0.05).

Participants who reported "waking too early" had earlier sleep onset and wake times compared to those without early awakenings (Figure 11). However, again their total sleep duration was similar to the non-early-waking group, indicating that while they woke earlier, they also went to bed earlier. This finding suggests that early waking may not necessarily indicate reduced sleep duration but rather an adjustment in sleep timing.

**Figure 11.**
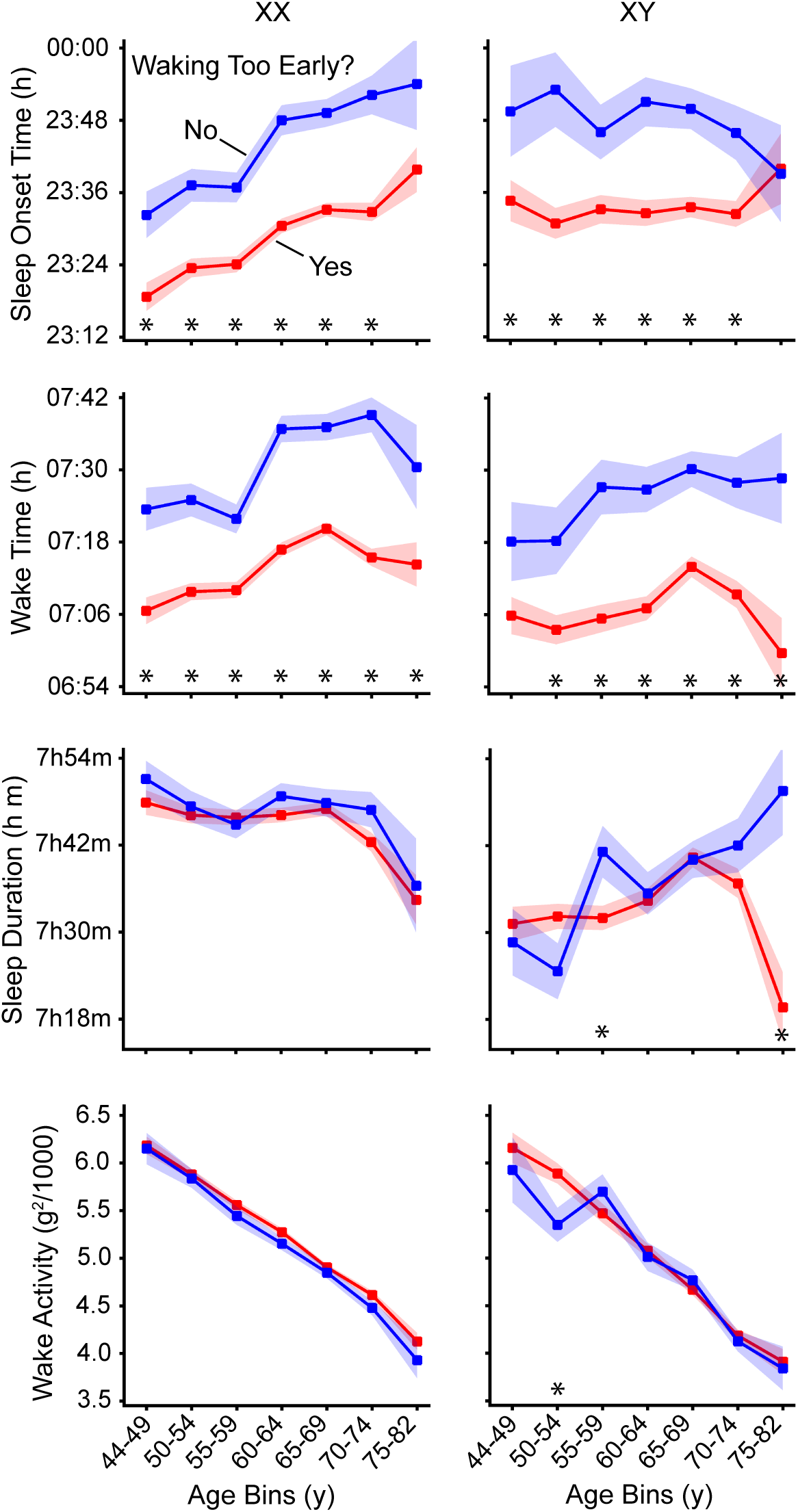
Sleep and activity parameters for individuals self-reporting "waking too early." Sleep parameters and daily activity means are shown for participants who responded "yes" or "no" to whether they were "waking too early." Those who reported waking early (red) and those who did not (blue) are shown with standard error bars. Those self-reporting waking too early go to sleep earlier, resulting in similar sleep duration to those responding no. Daily wake activity levels are comparable between groups. Significance was tested using a two-sample t-test (* = p < 0.05).

### Objective Sleep and Activity Differ in Relation to Self-Reported Mood

Participants who reported recent depression (Figure 12) or anhedonia (Figure 13) exhibited consistently lower daily wake activity compared to those without these symptoms, across ages and both sexes. Recently depressed older women also showed reduced sleep duration, though this difference was not always present. No significant duration relationships were observed before the age of 54. An association between late sleep onset and depressive symptoms was particularly pronounced in men, underscoring the relevance of sleep timing for mental health.

**Figure 12.**
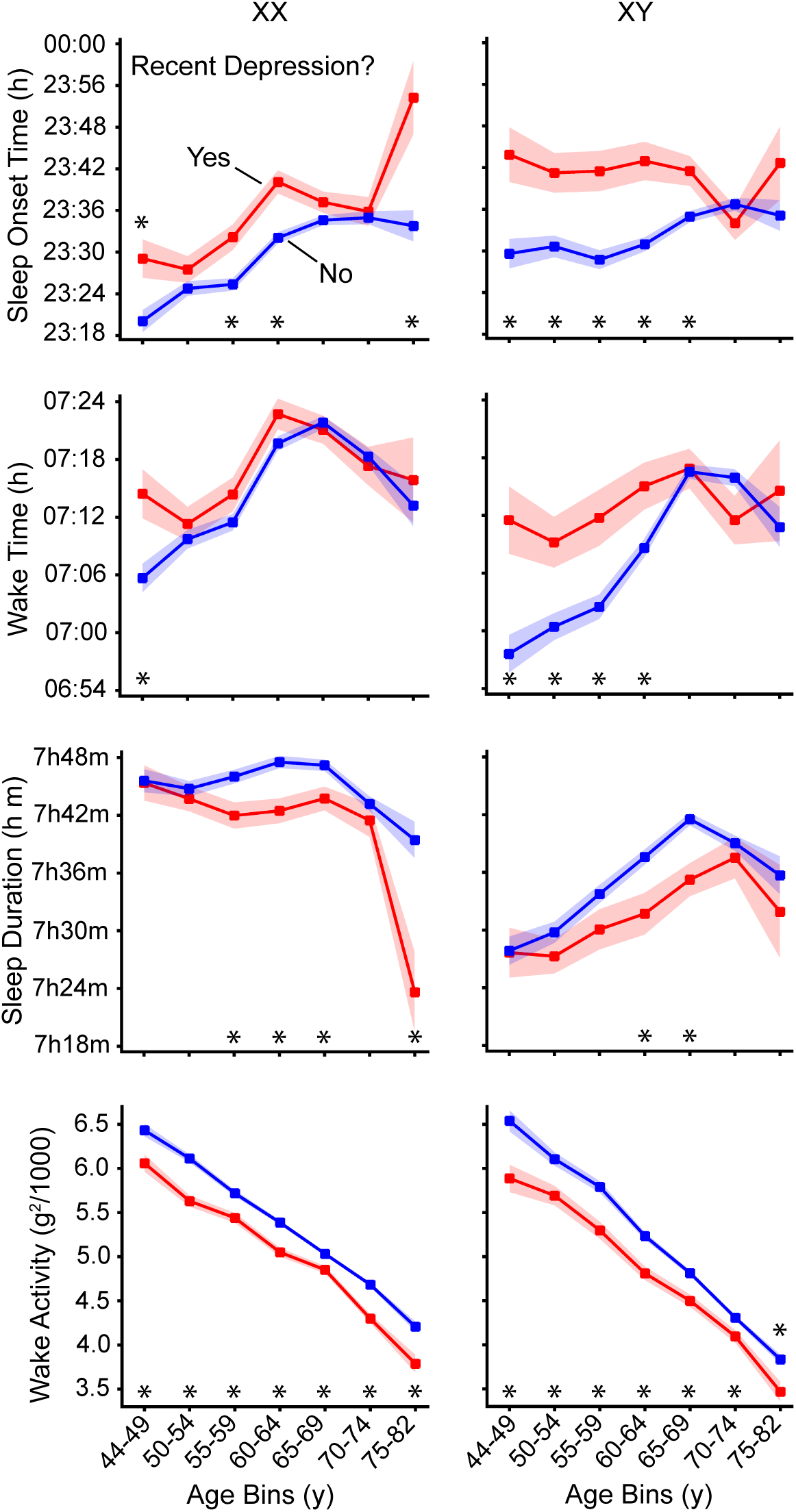
Sleep and activity parameters for individuals with recent self-reported symptoms of depression. Sleep parameters and mean wake activity are shown for participants responding "no" (not at all) or "yes" (any frequency) to feeling down, depressed, or hopeless over the prior two weeks. Individuals with recent depressive symptoms (red) and without (blue) are shown with standard error bars. Participants with symptoms of depression go to sleep slightly later with slightly shorter sleep duration. Symptoms of depression are associated with notably lower wake activity compared to other participants across ages. Significance was tested using a two-sample t-test (* = p < 0.05).

**Figure 13.**
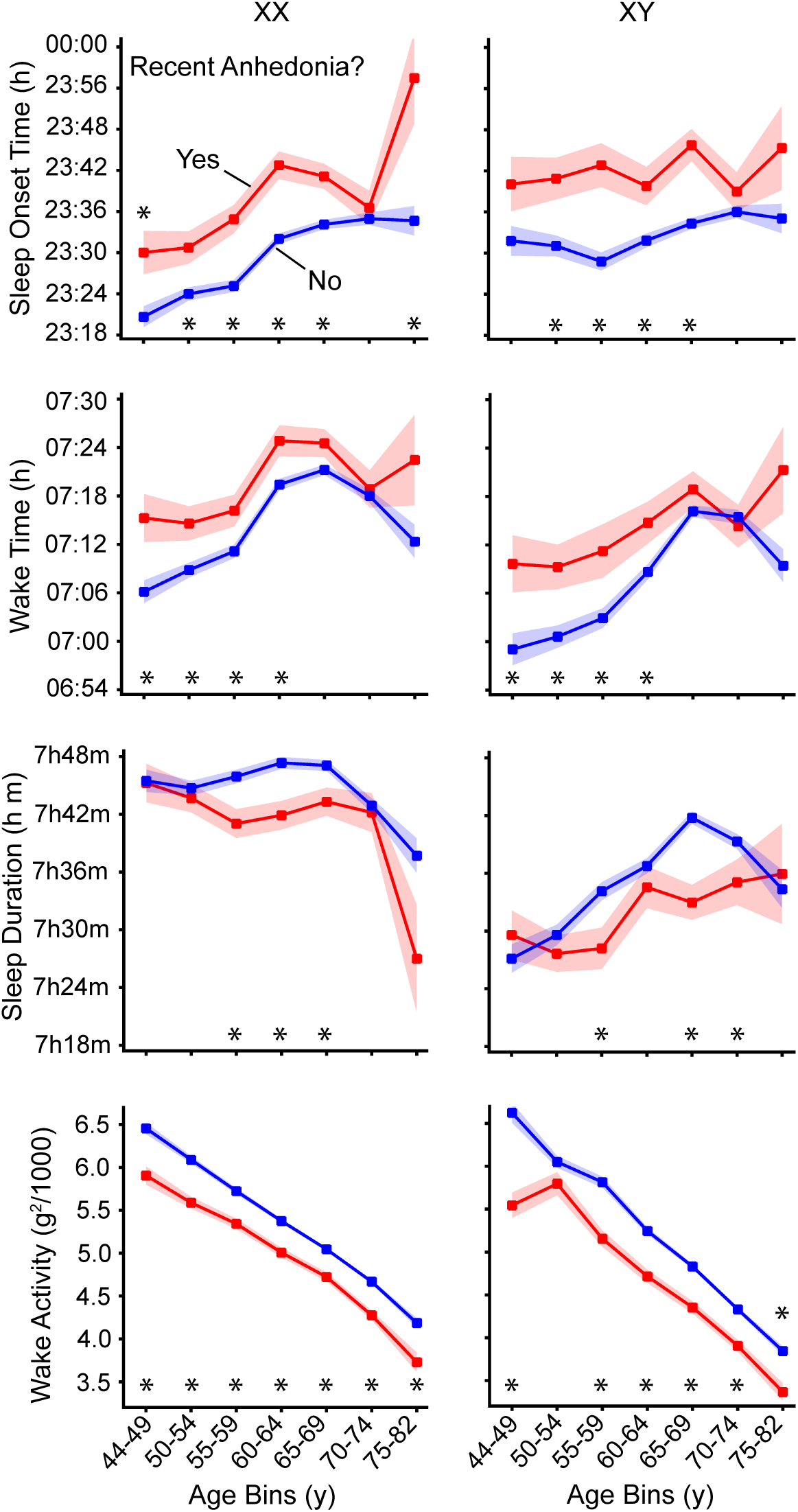
Sleep and activity parameters for individuals with recent self-reported anhedonia. Sleep parameters and daily activity means are shown for participants who responded "no" (not at all) or "yes" (any frequency) to experiencing little interest or pleasure in activities over the prior two weeks. Individuals with anhedonia (red) and without (blue) are shown with standard error bars. Participants with anhedonia go to sleep later and have slightly shorter sleep duration. The group reporting anhedonia exhibits lower wake activity compared to other individuals across all ages. Significance was tested using a two-sample t-test (* = p < 0.05).

## Discussion

Age- and sex-related differences in sleep were found and replicated across a large sample of 77,093 participants ages 44 to 82 years from the UK Biobank. Through rigorous quality control of actigraphy data, we identified complex patterns in sleep onset, wake time, and sleep duration that suggest reliable non-linear interactions between age and sex. Of particular interest are differences between women and men that attenuate in the oldest individuals. We also observed shifts in sleep and wake time linked to self-reported sleep behavior that did not associate with differences in overall sleep duration. By making these observations in a large sample with confidence, we contribute to a growing literature on sleep, aging, and mental health. The complex relations between self-reported sleep and the objective measures further underscore the importance of objective sleep measures in understanding age- and sex-specific sleep patterns.

### Sleep Patterns Reveal Reliable Non-Linear Interactions with Age and Sex

Men exhibited shorter sleep durations compared to women, especially in the middle-aged groups, where men tended to go to sleep later and wake up earlier than women. This pattern aligns with previous findings that suggest men sleep less which could be due to occupational, social, or biological factors (Meadows *et al*., 2008; Burgard and Ailshire, 2013). Of interest to study of aging, in older men and women, this sleep disparity narrowed, with both sexes achieving more similar sleep durations. Specifically, men’s sleep duration was increased in older age, except in the very oldest individuals, while women’s sleep duration was higher but then shortened in the eighth decade (Figure 4), a finding consistent with research that shows age-related differences in sleep are more pronounced in men (Sterniczuk and Rusak, 2017).

### Sleep Patterns Differ Across the Week in an Age-Dependent Manner

For individuals under 60, significant differences in sleep timing were observed between weekdays and weekends, with later sleep onset and wake times on weekends (Figure 5). This robust effect has multiple possible causes. One possible explanation for this pattern is social jetlag, a phenomenon in which individuals accumulate sleep debt during the workweek due to work or social obligations and then compensate by sleeping longer on weekends (Wittmann *et al*., 2006; Saxvig, Bjorvatn and Waage, 2023). Another possibility is that these patterns reflect a preference for sleeping longer on weekends not solely as a means of recovering sleep debt but as a form of leisure or enjoyment (Horne, 2016). In our study, men exhibited less pronounced weekend differences in terms of sleep onset, as their sleep onset times were consistently later even on weekdays. This behavior might indicate a prioritization of nighttime leisure activities over sleep, aligning with previous findings that link late bedtimes to recreational or social preferences (Burgard and Ailshire, 2013).

Future work will be required to determine the mechanisms behind these clear and reliable shifts in sleep onset and wake times that differ across the week. The presence of sleep differences during the week in younger populations raises concerns about its potential long-term effects on health, as chronic sleep deprivation has been linked to a variety of health problems, including metabolic disorders and cardiovascular disease (Wittmann *et al*., 2006; Roenneberg *et al*., 2019) although the relation of sleep duration and health consequences is nuanced (Fjell and Walhovd, 2024).

Reflecting a robust age difference, older participants (those over 65 years) showed largely uniform sleep patterns throughout the week, possibly because their sleep is less constrained by obligations such as work that vary across days of the week. This reduction in the weekday / weekend gap with age has been observed in previous studies, where older adults tend to exhibit more stable sleep schedules due to retirement or fewer external time pressures (Kitamura *et al*., 2016). What is striking in the present data is the contrast of the weekly sleep patterns between the middle-aged and the oldest individuals (Figure 5).

### Self-Reported Sleep and Mental Health Associate with Objective Sleep Patterns

Self-reported sleep and mental health symptoms, including depression and anhedonia, were reliably associated with objective sleep parameters. As a first result, speaking to the validity of self-report measures, we initially explored and found that self-reported sleep duration parametrically predicted objectively measured sleep duration. While expected the consistency of the pattern across 14 independent groups of participants each revealing a parametric relation among all four levels of the self-report variable was nonetheless impressive (Figure 9).

Specifically, individuals self-reporting shorter sleep duration objectively slept less than those individuals reporting longer sleep duration. Revealing a difference between self-report and objective measures, the relation was compressed in that subjective estimates of short versus long sleep duration were considerably larger than the objectively measured differences, but the relationship was robust and existed for both men and women across all age group. These findings bolster confidence in the use of self-report to assess sleep behaviors, but also raise the nuance that subjective impressions may be systematically shifted relative to objective measures or consider other features of sleep (e.g., sleep quality or time trying to fall asleep) (Kushida *et al*., 2001; Bakken *et al*., 2014). The compression of differences may potentially also result from lower reliability of the self-reported durations. Additionally, differences in the distribution of self-reported sleep durations by sex and age help explain why younger men exhibit shorter objective sleep durations; a larger proportion of younger men reported shorter sleep (e.g., 6 hours), while reports of longer sleep (e.g., 8–9 hours) became more common with age—a shift not observed in women.

A further interesting finding was that self-report of ‘waking too early” was consistently associated with earlier objective wake times, as well as proportionate shifts in evening sleep onset times (Figure 11). As a result, total sleep duration was broadly similar regardless of self-report of wake time. This result is consistent with the self-report, which here focused specifically on wake time, but does indicate that endorsing “waking too early” is not a proxy for total sleep duration. Even more striking was that endorsing “too much sleep,’ a question that on the surface targets a construct expected to relate to sleep duration, was also not associated with an objective difference in total sleep duration. Rather, like endorsing early rising, self-report of “too much sleep” was associated with a shift in sleep-wake times with those reporting sleeping too much falling asleep and waking later than those who did not make such an endorsement. Furthermore, endorsing “too much sleep” was consistently linked to lower wake activity in both sexes across almost all age groups. Such an effect on activity was absent in those endorsing “waking too early”. These results indicate that self-reported sleep measures are capturing reliable differences in objective sleep parameters, but not in a simple or straightforward manner that could be intuited without the available objective sleep measures.

Of most interest, individuals reporting recent depression (Figure 12) or anhedonia (Figure 13) exhibited later sleep onset times and shorter sleep durations compared to those who did not endorse depressive symptoms. The effect was not observed in every group but was present and independently significant (replicated) in multiple groups across the age range including both sexes. These findings support the established link between sleep disturbances and depression (Hu *et al*., 2022). Interestingly, while depression was associated with reduced sleep duration, it was also linked to reduced wake activity, further suggesting that individuals with low mood may engage in less physical activity throughout the day, a pattern consistent with previous studies (Prince *et al*., 2017). A bidirectional relationship between sleep and depression is well-documented. Changes in sleep duration or timing can exacerbate mood disorders, while depression can lead to changes in sleep duration and timing (Joo *et al*., 2022). Our results corroborate the complex interaction between sleep and mood, particularly the association between sleep timing and mental health symptoms. Individuals with late sleep onset and late wake times, especially men, were more likely to report experiencing symptoms of depression, emphasizing the possible association of chronotypes with mental health. Prior research has shown that earlier sleep times are associated with a lower risk of depression, potentially due to greater exposure to daylight, which has mood-enhancing effects (Daghlas *et al*., 2021). While our results do not resolve the mechanism of the association between sleep parameters and symptoms of depression, the results illustrate that the relationship is present across many cohorts that vary by sex and age.

### Caveats and Limitations

While actigraphy is a valuable tool for large-scale, real-world sleep assessment, it does not provide the detailed sleep stage information available through PSG. PSG, as the gold standard for identifying specific sleep disorders and sleep stages, is not well-suited for large-scale studies due to the burden and cost. Further, PSG necessarily disturbs normal sleep patterns and is not suited to measure natural variations in sleep duration. Advances in actigraphy technology, including the integration of heart rate and heart rate variability measurements, may enhance their ability to estimate sleep stages. Ambulatory PSG may also become viable at scale in the future.

Additionally, the UK Biobank cohort includes cross-sectional participants ages 44 and older, limiting the ability to the chart patterns in younger adults and to explicitly estimate how sleep patterns change with age. Further, ethnic variation among UK Biobank participants is limited, and different sample compositions or samples from different societies might yield other results (Willoughby *et al*., 2023). Early sleep disturbances in younger cohorts might be critical predictors of mental health disorders or neurodegenerative conditions. Cohort effects, such as the possibility of differential survivorship in the oldest groups, could be clarified by a longitudinal design. Future research should extend these analyses to younger age groups and longitudinal designs to investigate how sleep patterns influence the onset of illness and changes in symptoms.

### Future Directions and Implications

In addition to the findings themselves, the present results underscore the importance of age- and sex-specific analyses to understand sleep patterns. An immediate implication of our results is that non-linear sleep patterns and interactions should be considered when exploring the relation of sleep and other factors. For example, regressing the linear effects of age or the age x sex interaction may be insufficient to capture the complex patterns observed here. The present normative description of age and sex differences on sleep patterns is thus valuable as a reference to guide future study designs and analyses that target older populations.

The variation in sleep patterns between groups and individuals within the groups were marked. Finding reproducible relations between self-reported perceptions of sleep and mood indicate that the variation contains structured signal not happenstance noise. However, the present results do not provide insights into the biological or other factors that cause the individual differences. Future research, including using the UK Biobank data itself, might explore integrating genetic factors and neuroimaging to examine how sleep patterns are influenced by genetic predispositions and associate with variations in brain structure. For instance, specific genetic polymorphisms related to circadian rhythm regulation and sleep architecture could interact with age- and sex-related changes in sleep (Archer *et al*., 2010). Combining actigraphy data with genotyping could potentially help identify genetic markers associated with sleep resilience or susceptibility to sleep disturbances, which may be predictive of cognitive decline (Dashti *et al*., 2019; Jones *et al*., 2019). The present normative characterization of sleep patterns and how they differ by age and sex provides a needed foundation for such future research.

A further intriguing possibility along these lines is that some of the genetic variations associated with sleep patterns or sleep quality in early life may be risk factors for brain aging and dementia later in life because they are linked to broad sequelae that affect the brain and body. That is, some of the genetic variation associated with dementia risk may be related to mechanistic links involving sleep that are associated with health throughout life and may indirectly reflect risk for cognitive decline.

Another future direction will be to explore the association of sleep with brain aging. Sleep pattern differences that vary by age and sex could inform structural brain differences and longitudinal change. Although prior neuroimaging studies have suggested links between poor sleep and brain atrophy, recent large-scale analyses challenge this view (Fjell *et al*., 2023), highlighting the need for future research to disentangle sleep-related correlates from causal effects on brain structure. Longitudinal studies incorporating structural MRI could be undertaken to examine how age-related sleep differences correlate with brain atrophy and effects on white matter integrity (Spira *et al*., 2016).

## Acknowledgements

The UK Biobank analyses were conducted under application Number 67237. This research was supported by NIA grants R01AG067420, R01DC014296, and P30AG062421, the Simons Foundation grant 811255, and the Vranos Family Fund. M.L.E. is supported by NIA grant K00AG068432. We thank Kevin Anderson for valuable assistance in handling the UK Biobank data and Timothy O’Keefe for neuroinformatic support.

